# Phosphoregulation of the Rad51 auxiliary factor Swi5-Sfr1

**DOI:** 10.1101/2023.03.23.533938

**Authors:** Pengtao Liang, Katie Lister, Luke Yates, Bilge Argunhan, Xiaodong Zhang

## Abstract

Homologous recombination (HR) is a major pathway for the repair of DNA double-strand breaks, the most severe form of DNA damage. The Rad51 protein is central to HR, but multiple auxiliary factors regulate its activity. The heterodimeric Swi5-Sfr1 complex is one such factor. It was previously shown that two sites within the intrinsically disordered domain of Sfr1 are critical for the interaction with Rad51. Here, we show that phosphorylation of five residues within this domain regulates the interaction of Swi5-Sfr1 with Rad51. Biochemical reconstitutions demonstrated that a phosphomimetic mutant version of Swi5-Sfr1 is defective in both the physical and functional interaction with Rad51. This translated to a defect in DNA repair, with the phosphomimetic mutant yeast strain phenocopying the previously established interaction mutant. Interestingly, a strain in which Sfr1 phosphorylation was blocked also displayed sensitivity to DNA damage. Taken together, we propose that controlled phosphorylation of Sfr1 is important for the role of Swi5-Sfr1 in promoting Rad51-dependent DNA repair.

## INTRODUCTION

DNA damage occurs through both exogenous and endogenous sources, making it an unavoidable occurrence in the life of all organisms. The arsenal of DNA repair processes is vast and varied, but a common feature is the use of the intact complementary DNA strand as a template for synthesis-dependent repair of the damaged strand. A single DNA double-strand break (DSB) interrupts the continuity of the DNA molecule, precluding the use of the complementary strand as a template. Taken together with the fact that a single DSB is sufficient to cause cell death, DSBs pose a particularly challenging threat to the cell. Homologous recombination (HR) is a major DSB repair mechanism that circumvents the problem of disrupted continuity by identifying another region in the genome that shares high sequence identity (i.e., homology) to the damaged DNA and utilizing this region as a template for synthesis-dependent repair (1).

The DNA ends at a DSB site are processed by nucleases to generate 3’-ended single-stranded DNA (ssDNA) tails that are first bound by the ssDNA-binding protein RPA (2). RPA is then replaced with the RecA-family recombinase Rad51, which oligomerizes to form a right-handed helical filament with the ssDNA running along its central axis (3). This nucleoprotein filament interrogates the genome to identify a homologous region of intact double-stranded DNA (dsDNA) and invades into the duplex DNA, displacing the identical strand and forming base-pairs with the complementary strand, leading to the formation of a key recombination intermediate known as a displacement loop (D-loop) (4, 5). The D-loop can be expanded by Rad51-driven strand-exchange and extension of the 3’-ended invading strand, which serves as a primer for DNA synthesis. The invading strand then dissociates from the duplex, and having been extended, can anneal to the ssDNA exposed on the other side of the DSB. Following gap filling by further DNA synthesis and ligation of resultant nicks, DSB repair by HR is complete. This model of HR is known as synthesis-dependent strand annealing (SDSA) and is strictly dependent on Rad51 (1). Thus, the formation/stabilisation of the Rad51 nucleoprotein filament is highly regulated.

The Swi5-Sfr1 (S5S1) auxiliary factor complex was first identified as a Rad51 regulator in the fission yeast *Schizosaccharomyces pombe* (6). Biochemical reconstitutions with purified proteins directly demonstrated that S5S1 physically interacts with Rad51 and stimulates its strand exchange activity by stabilizing Rad51 nucleoprotein filaments (7, 8). Homologues have since been identified in mice and humans (9, 10), and experiments with mouse proteins indicated that the mechanisms underlying Rad51 potentiation by S5S1 are largely conserved (11–14).

Structural analysis of *S. pombe* S5S1 demonstrated that it is an elongated heterodimer: the crystal structure of the C-terminal half of Sfr1 in complex with Swi5 (S5S1C) revealed that this truncated complex forms a coiled-coil, whereas the N-terminal half of Sfr1 (Sfr1N) was shown to be intrinsically disordered by circular dichroism spectroscopy (CD) and nuclear magnetic resonance (NMR) analysis (15–17). These two structurally distinct domains were also found to comprise disparate functional modules (16). While S5S1C could activate Rad51, ∼10-fold more protein was required than wild type for maximal stimulation, and a S5S1C-Rad51 complex could not be detected by immunoprecipitation, suggesting that the physical interaction is very weak/transient. By contrast, Sfr1N was found to co-immunoprecipitate with Rad51, but did not stimulate Rad51 activity, leading the authors to propose that Sfr1N serves as an anchor to tether S5S1C to activate Rad51 (16). NMR interaction analysis identified two sites in Sfr1N that bind Rad51 (17): Site 1 (Ser84–Thr114) and Site 2 (Thr152-Ser168). Mutation of seven Lys/Arg residues in these sites to Ala (S5S1-7A) severely impaired the physical and functional interaction between S5S1 and Rad51 (17).

Although the underlying molecular mechanisms are only just being elucidated, the regulation of HR by auxiliary factors has long been appreciated (18). By contrast, the role of post-translational modifications (PTMs), which represent an additional layer of regulation, is poorly understood. Although modern proteomics studies have identified many PTMs of HR auxiliary factors, how such modifications affect HR is not clear, despite increasing evidence suggesting that they are important (19). Phosphoproteomics identified multiple phosphorylated residues located in Sfr1N (20–24): S24, T73, S109, S116, S165. Interestingly, several of these are situated in/around the Rad51 interaction sites.

Here, we investigated the possibility that phosphorylation of Sfr1 regulates the interaction of S5S1 with Rad51. Mutation of the five S/T residues to phosphomimetic D residues severely impaired the binding of S5S1 to Rad51. Furthermore, the stabilisation of Rad51 filaments by S5S1 was compromised by the phosphomimetic mutations, and this translated to a substantial reduction in the stimulation of Rad51-driven strand exchange. Importantly, these defects were comparable in magnitude to the S5S1-7A mutant, a bonafide interaction mutant (17). Yeast strains expressing phosphomimetic mutants of Sfr1 phenocopy *sfr1-7A*, displaying the same DNA damage sensitivity as *sfr1Δ* only in the absence of the Rad55-Rad57 auxiliary factor complex. Interestingly, cells expressing a non-phosphorylatable version of Sfr1 (five S/T to A mutations) also displayed DNA damage sensitivity, despite this mutant protein being biochemically comparable to wild type. Taken together, we propose that phosphorylation negatively regulates the interaction of S5S1 with Rad51, providing a means to fine-tune Rad51 potentiation by S5S1.

## RESULTS

### Biochemical characterisation of S5S1 mutant variants

To investigate if phosphorylation of Sfr1 is pertinent to the function of S5S1 in HR, we mutated several phosphorylation sites of Sfr1 and purified the protein in complex with Swi5 **(Figs. 1A,B)**. Since the proteins were purified from *Escherichia coli*, wild-type S5S1 serves as the unphosphorylated sample. A variant in which five S/T residues were mutated to the phosphomimetic D residue serves as a proxy for the phosphorylated sample (S5S1-PM, <u>p</u>hospho<u>m</u>imetic). Given that several of these S/T residues are situated in/around the two Rad51 interaction sites within Sfr1N (17) **(Fig. 1A)**, any defects observed for S5S1-PM could be explained by the loss of S/T residues that are important for the interaction with Rad51. To control for this, we also purified a variant in which the S/T residues of interest were mutated to A (S5S1-NON, <u>non</u>-phosphorylatable). Finally, a previously established Rad51 interaction mutant was also purified for comparison (17) (S5S1-7A, seven K/R residues mutated to A). Consistent with the fact that the C-terminal half of Sfr1 is important for complex formation with Swi5 (16), the oligomeric state of the S5S1 heterodimer was not affected by any of these mutations in the N-terminal domain **(Fig. 1C)**.

**Figure 1.**
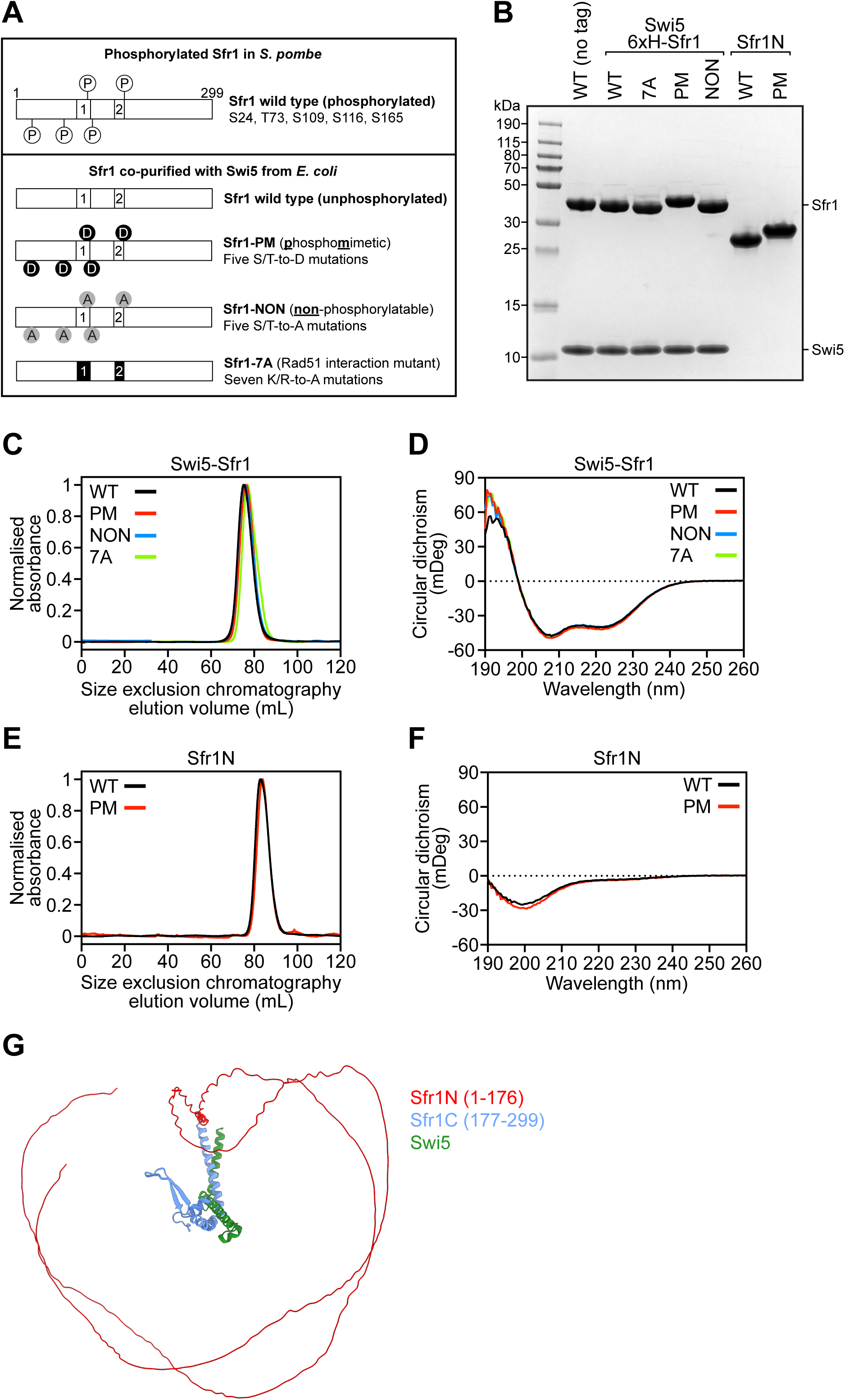
Biochemical characterisation of Swi5-Sfr1 (S5S1) mutants. **(A)** Schematic of Sfr1 mutants. **(B)** SDS-PAGE analysis of purified S5S1 complexes and Sfr1N variants. 2 µg of each protein was loaded. **(C,E)** Size-exclusion chromatography analysis of indicated proteins. **(D,F)** Circular dichroism spectra of indicated proteins. **(G)** AlphaFold 2 Multimer models of S5S1 and S5S1-PM overlayed with Sfr1C as a reference point. Sfr1N, red. Sfr1C, blue. Swi5, green. The prediction suggests that Sfr1N remains disordered irrespective of the phosphomimetic mutations.

S/T residues located in intrinsically disordered domains are commonly targeted for phosphorylation, and in some cases, these phosphorylation events promote folding into structured domains (25, 26). To examine whether the phosphorylation of Sfr1 might alter its structure, CD spectroscopy was employed to obtain a course readout of global S5S1 structure. The CD spectrum of S5S1 resembled an average of disordered and α-helical signals **(Fig. 1D)** (27), as expected from the disordered Sfr1N and predominantly coiled-coil nature of S5S1C (16, 17). The spectra of other mutants examined here were essentially identical, suggesting that the introduction of phosphomimetic mutations does not cause any obvious change in the structural landscape of S5S1. However, the presence of S5S1C could mask any subtle changes in the disordered state of Sfr1N. We therefore purified Sfr1N and a version containing the phosphomimetic mutations (Sfr1N-PM) **(Fig. 1B)**. Sfr1N and Sfr1N-PM both show similar elution profiles from size exclusion chromatography **(Fig. 1E)** and their CD spectra were characteristic of a disordered protein (27) **(Fig. 1F)**, as was previously observed for wild-type Sfr1N (17). These experimental data are consistent with predicted structural models of both S5S1 and S5S1-PM generated by AlphaFold 2 multimer (28), which show that Sfr1N of both complexes is expected to be completely disordered, except for a short α-helix at the extreme C-terminus of the disordered domain (P166-C170) **(Fig. 1G)**. Taken together, these results indicate that the global structure of S5S1 or the disordered state of Sfr1N are not obviously altered by the introduction of phosphomimetic mutations in Sfr1N.

### Phosphomimetic mutations in Sfr1 impair the binding of S5S1 to Rad51

Next, we sought to directly test whether the binding of Rad51 to S5S1 is affected by the phosphomimetic mutations in Sfr1. To this end, the hexahistidine tag at the N-terminus of Sfr1 was utilised in pull-down assays. As expected, when purified Rad51 **(Fig. S1)** was mixed with S5S1 and complexes were precipitated using nickel resin, a significant amount of Rad51 was seen to co-precipitate with S5S1 **(Fig. 2A)**. By contrast, substantially less Rad51 was observed when S5S1-7A was precipitated. Strikingly, a similar reduction in the co-precipitation of Rad51 was seen when S5S1-PM was employed, and this was not the case for S5S1-NON.

**Figure 2.**
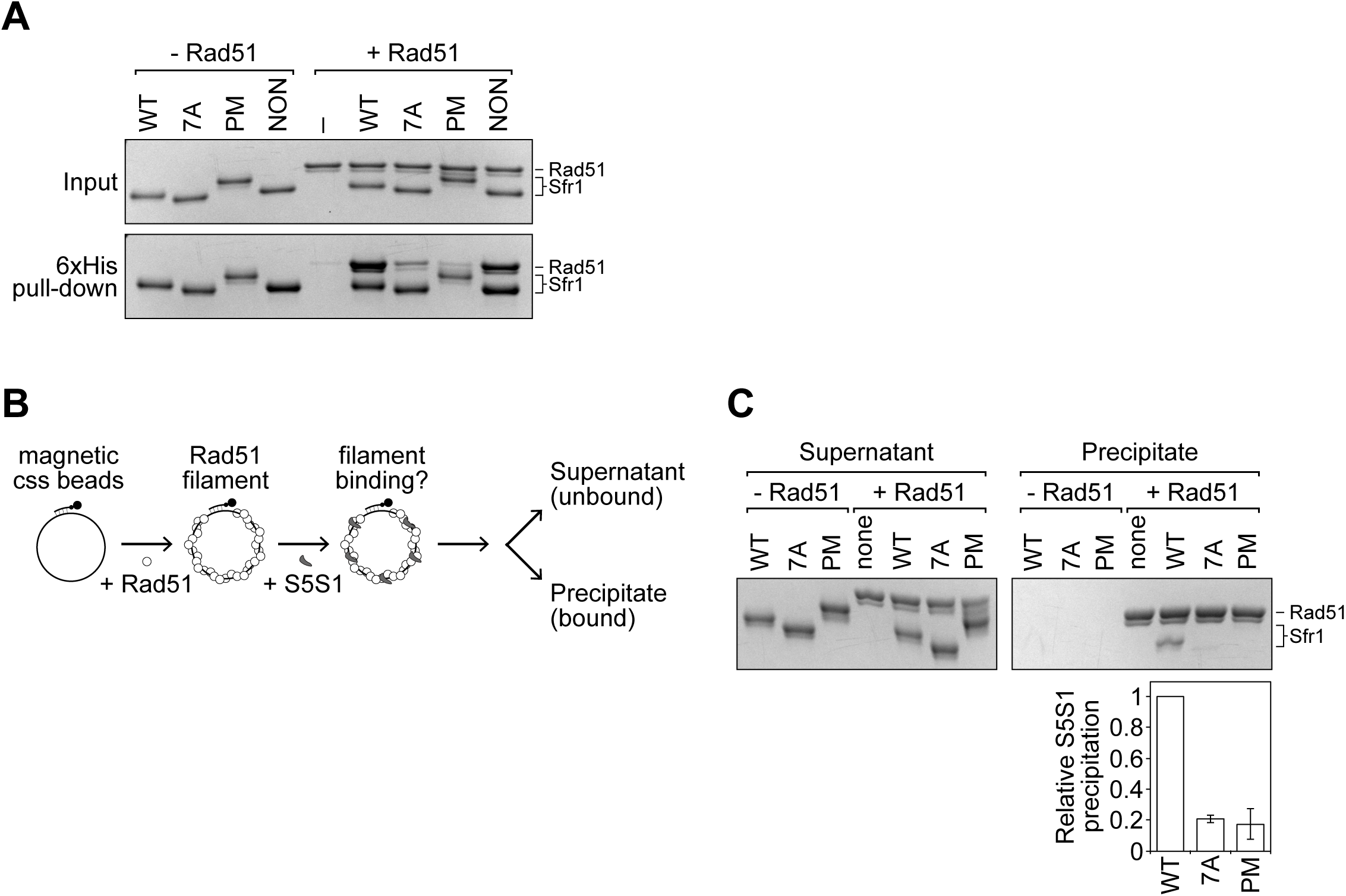
The physical binding of S5S1 to Rad51 is impaired by phosphomimetic mutations in Sfr1. **(A)** Pull-down experiments with nickel resin were performed with purified Rad51 and the indicated variant of hexahistidine-tagged S5S1 to examine the interaction in-solution (500 nM each protein). The content of precipitated resin from each reaction was examined by SDS-PAGE and Coomassie staining. –, protein omitted. **(B)** Schematic of the filament interaction assay. **(C)** Rad51 filaments were pulled-down after incubation with the indicated variant of S5S1. The amount of Rad51 and Sfr1 that was precipitated was determined by SDS-PAGE and Coomassie staining. Sfr1 signal was normalised to Rad51 and expressed relative to wild type. Averages are plotted. n = 3, error bars represent standard deviation.

Early biochemical analyses suggested that S5S1 stabilises Rad51 filaments but does not function as a canonical mediator that directly promotes Rad51 assembly on ssDNA (e.g., Rad52 in yeast and BRCA2 in humans) (8). This was largely corroborated by more recent single-molecule experiments demonstrating that S5S1 reduces the dissociation rate of Rad51-ssDNA complexes without affecting the association rate i.e., S5S1 stabilises Rad51 filaments but does not promote filament formation per se (29). We interpret these results to mean that the physiological substrate S5S1 acts on is likely to be the Rad51-ssDNA filament rather than free Rad51. With this in mind, we employed an assay to test the binding of S5S1 to Rad51-ssDNA filaments **(Fig. 2B)**. Briefly, Rad51 filaments were formed on ssDNA that was annealed to a biotinylated oligonucleotide, and reactions were supplemented with S5S1. Following a short incubation, the Rad51-ssDNA complexes (precipitate) were separated from the unbound proteins (supernatant) using streptavidin-coated magnetic beads, and both fractions were analysed by SDS-PAGE. Whereas a clear Sfr1 band was observed for S5S1, the Sfr1 band for S5S1-PM was barely detectable, corresponding to an approximately five-fold reduction in the co-precipitation of S5S1 with Rad51 **(Fig. 2C)**. A similar reduction was observed for S5S1-7A, the established Rad51 interaction mutant (17). These results suggest that phosphorylation of Sfr1 at these five residues severely disrupts the interaction of S5S1 with Rad51.

### The functional interaction of S5S1 with Rad51 is compromised by phosphomimetic mutations in Sfr1

Having determined that the physical binding of Rad51 to S5S1-PM is severely impaired, we next asked whether the magnitude of the binding defect is functionally consequential. A critical mechanism through which S5S1 potentiates Rad51 is by stabilising Rad51-ssDNA filaments (8, 29). Although S5S1C possesses this activity, abrogation of the binding to Rad51 by mutating the interaction sites in Sfr1N (i.e., S5S1-7A) severely impaired the efficiency of filament stabilisation, with several-fold more protein required to effectively stabilise filaments (16, 17). To test whether S5S1-PM can effectively stabilise Rad51-ssDNA filaments, we examined filaments by negative-stain electron microscopy (NSEM). The formation of human RAD51-ssDNA filaments in the presence of ATP-Mg^2+^ lead to the visualisation of heterogenous species by NSEM, including short, ordered filaments, disordered filaments, and ring-like oligomers (30). We observed similar species with *S. pombe* Rad51 filaments (116-mer ssDNA) formed in the presence of ATP-Mg^2+^ **(Fig. 3A)**. We reasoned that, under these conditions, stabilisation of Rad51-ssDNA filaments by S5S1 would result in increased filament length. Indeed, longer filaments were observed upon inclusion of S5S1, and there was a concomitant reduction in other Rad51 species, indicative of improved filament stability **(Fig. 3B)**. The addition of either S5S1-PM or S5S1-7A seemingly resulted in an intermediate state, with some longer filaments observed **(Fig. 3C,D)**. These qualitative observations were corroborated by measurements of filament length **(Fig. 3E)**, which revealed that, while S5S1-PM and S5S1-7A were able to partially stabilise Rad51 filaments, they were unable to do so to the same extent as wild-type S5S1. These differences in filament stabilisation were statistically significant, and the defect of S5S1-PM was comparable to S5S1-7A.

**Figure 3.**
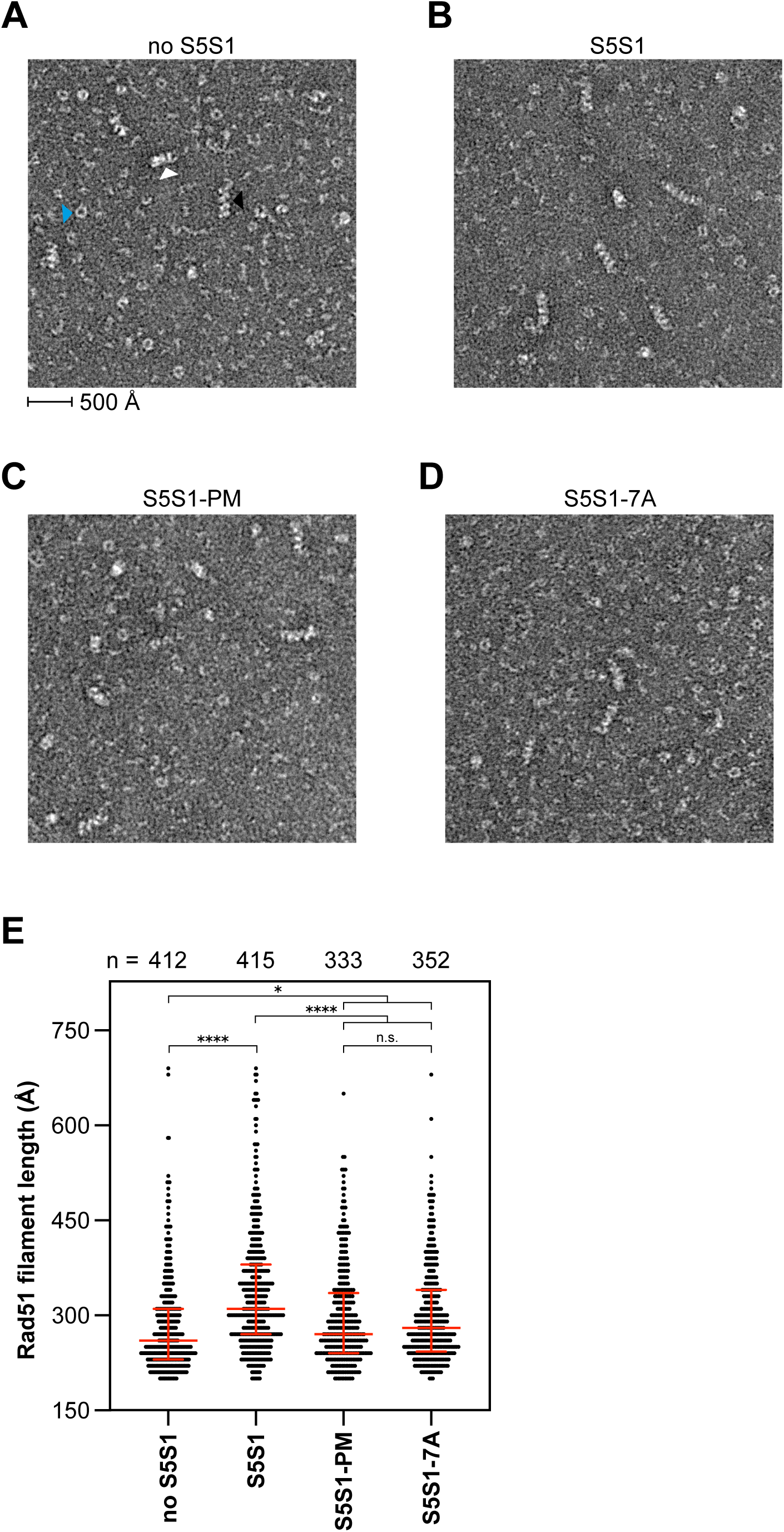
S5S1-PM is impaired for Rad51 filament stabilisation. Representative electron micrographs of Rad51 filaments formed on 116-mer ssDNA in the absence of S5S1 **(A)**, or in the presence of S5S1 **(B)**, S5S1-PM **(C)**, or S5S1-7A **(D)**. Different Rad51 species are highlighted in **(A)**: white arrowhead, disordered filament; black arrowhead, ordered filament; blue arrowhead, ring-like oligomer. **(E)** Quantification of filament length from reactions supplemented with the indicated S5S1 variant. Red bars represent median and interquartile range. Number of filaments quantified per sample (n) is shown above the graph. One-way ANOVA with multiple comparisons correction (Tukey) was conducted to test statistical differences. *, *P* < 0.05. ****, *P* < 0.0001. n.s., not significant (*P* = 0.9958).

Next, we sought to examine whether the defect of S5S1-PM in filament stabilisation translated to an impairment in the stimulation of Rad51-driven DNA strand exchange. A strand exchange assay with plasmid-sized DNA substrates was employed to directly test this **(Fig. 4A)**. Briefly, the Rad51 filament is formed on circular ssDNA (cssDNA) and homologous linear dsDNA (ldsDNA) is added. Rad51 drives the pairing of cssDNA and ldsDNA, forming joint molecules (JMs), which are the reaction intermediates. Following strand transfer over the length of the DNA, linear ssDNA and nicked circular DNA (NC) are generated as the reaction products. This assay requires the eukaryotic ssDNA-binding protein Replication Protein A (RPA) (7), which was also purified to homogeneity **(Fig. S1)**.

**Figure 4.**
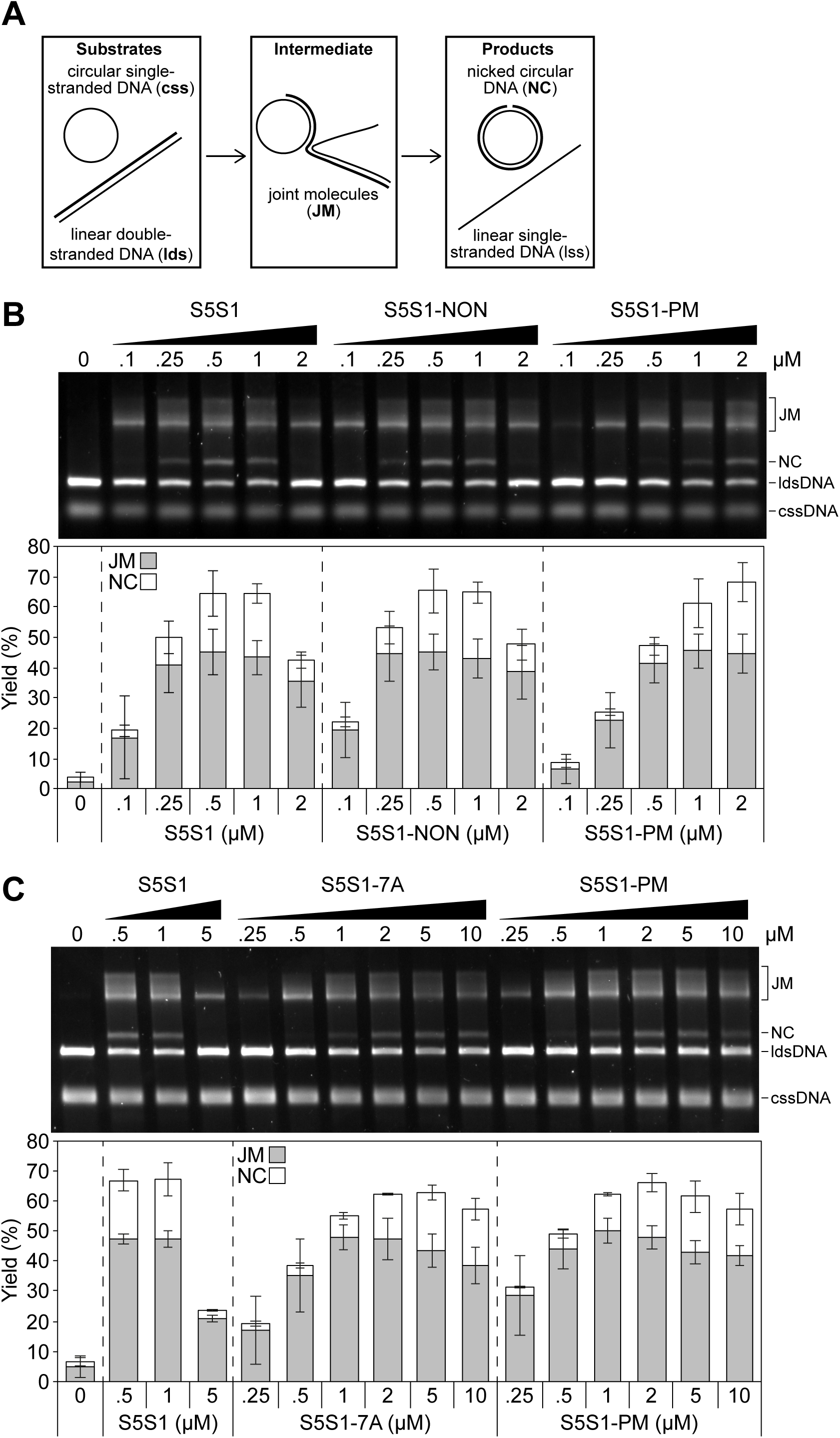
Phosphomimetic mutations in Sfr1 impair the stimulation of Rad51-driven DNA strand exchange by S5S1. **(A)** Schematic of the strand exchange assay. **(B,C)** All incubations were at 37°C. Rad51 (5 µM) was incubated with cssDNA (10 µMnt) then supplemented with the indicated concentration of a S5S1 variant. RPA (1 µM) was then added, and following a further incubation, reactions were initiated through the addition of ldsDNA (10 µMnt). After a 2 h incubation, reactions were deproteinised, DNA species were resolved by agarose gel electrophoresis and visualised. Percentage of JM (reaction intermediate) and NC (reaction products) were quantified, and averages were plotted. n = 6 for **(B)**, n = 3 for **(C)**, error bars represent standard deviation.

We first confirmed that the hexahistidine tag introduced at the N-terminus of Sfr1 did not affect the ability of S5S1 to stimulate Rad51-driven strand exchange **(Fig. S1)**. In the absence of S5S1, virtually no JM or NC was observed, indicating that Rad51 is unable to drive homologous DNA pairing and subsequent strand exchange under these conditions **(Fig. 4B)**. The inclusion of sub-stoichiometric amounts of S5S1 strongly stimulated Rad51 activity, with peak stimulation observed at around 0.5 µM (S5S1:Rad51 of 1:10). This stimulatory effect was diminished at higher concentrations, with significantly less NC observed at 2 µM (and even less at stoichiometric concentrations, see next paragraph). This is consistent with previous observations (7, 8, 17). Strikingly, the stimulation of Rad51 by S5S1-PM was severely impaired, with ≥2 µM protein required to achieve stimulation comparable to that of wild type at 0.5 µM **(Fig. 4B)**. Given that S5S1-NON showed a stimulation profile that is indistinguishable from wild type, these results strongly suggest that the introduction of phosphomimetic mutations rather than the loss of S/T residues in Sfr1 compromises the stimulation of Rad51 by S5S1.

Since the results with S5S1-PM are reminiscent of previous observations with S5S1-7A, the established Rad51 interaction mutation, the two mutants were compared to each other side-by-side over a broader concentration range. Consistent with previous observations, S5S1-7A showed peak Rad51 stimulation at 2–5 µM, with only a marginal reduction in stimulation at 10 µM; this is in stark contrast to wild type, where a substantial loss of stimulation was seen even at 5 µM **(Fig. 4C)**. Overall, the magnitude of the defects observed for S5S1-PM are very similar to S5S1-7A. Taken together, these results suggest that phosphorylation of Sfr1 at these five residues significantly impairs the ability of S5S1 to stabilise Rad51-ssDNA filaments and stimulate Rad51-driven DNA strand exchange.

In addition to an impairment in the interaction with Rad51, S5S1-7A was previously shown to have severe defects in DNA binding, although, based on multiple lines of indirect evidence, the authors argued that this DNA binding activity may be impertinent to the role of S5S1 in DNA repair (17). Nevertheless, we examined the ability of S5S1-PM to bind DNA in electrophoretic mobility shift assays. Unlike S5S1-7A, S5S1-PM was able to shift both ssDNA **(Fig. S2)** and dsDNA **(Fig. S2)** significantly, although not to the same extent as wild-type S5S1. Importantly, S5S1-NON shifted DNA comparably to wild type across all concentrations examined, suggesting that the introduction of phosphomimetic residues mildly impairs DNA binding. Despite the clear difference in DNA binding between S5S1-7A and S5S1-PM, the fact that both mutants are similarly defective for filament stabilisation **(Fig. 3E)** and strand exchange stimulation **(Fig. 4C)** is consistent with the notion that the DNA binding activity of S5S1 is not important for Rad51 potentiation (17).

### The phosphorylation status of Sfr1 is important for DNA repair

Our biochemical analyses suggested that phosphorylation of Sfr1 severely impairs the S5S1-Rad51 interaction. However, it remained unclear whether this constitutes a physiological mechanism to regulate recombinational DNA repair. To test this, we constructed *S. pombe* strains in which the native *sfr1^+^* gene was replaced with an allele encoding phosphomimetic D substitutions [*sfr1-PM(D)*] or E substitutions [*sfr1-PM(E)*] at the five S/T residues of interest. A strain encoding the non-phosphorylatable mutant (*sfr1-NON*) was also constructed. These strains were then assayed for DNA damage sensitivity via a standard spot-test. *S. pombe* possesses an alternative ultraviolet light (UV) damage repair pathway that requires HR for the repair of endonuclease-induced DNA breaks; the response to UV damage is therefore a suitable measure of HR proficiency (31). In contrast to the strain in which the *sfr1^+^* gene was deleted (*sfr1Δ*), the phosphomutants showed normal or near-normal growth following acute exposure to UV, suggesting that DNA repair is not obviously perturbed in these strains **(Fig. 5A)**.

**Figure 5.**
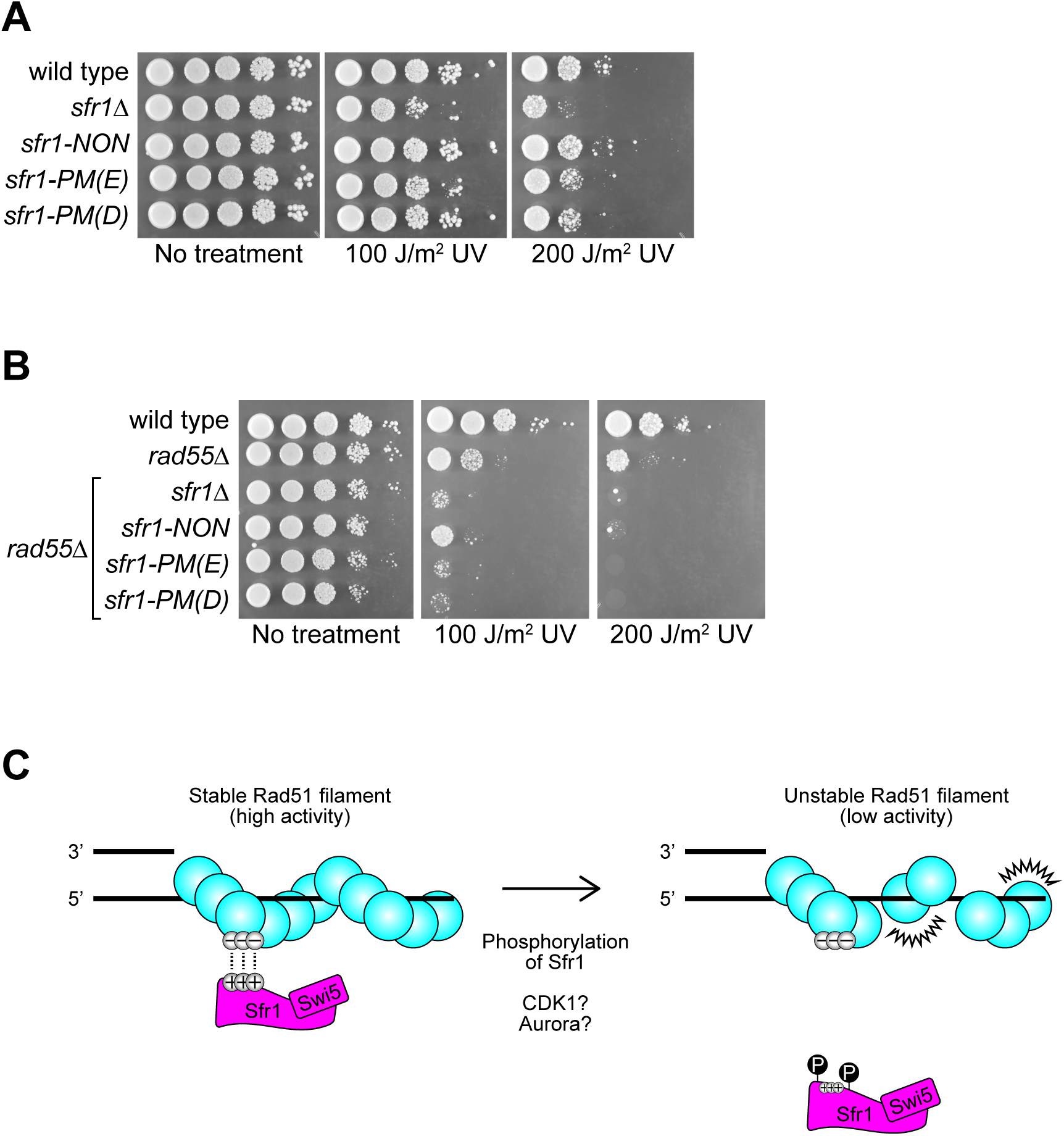
The phosphorylation status of Sfr1 is pertinent for the role of S5S1 in promoting Rad51-dependent DNA repair. **(A, B)** Log phase cultures of the indicated *S. pombe* strains were serially diluted (10-fold) and spotted onto solid rich media. Plates were not treated (control) or subjected to the indicated dose of acute UV irradiation. Plates were imaged following growth at 30°C for 3– 4 days. **(C)** Working model depicting how S5S1 binds to Rad51 and promotes DNA repair.

Given the biochemical defects observed for S5S1-PM, this result may seem surprising. However, through examination of the *sfr1-7A* mutant, it was previously shown that Rad55-Rad57 can suppress defects in the interaction of S5S1 with Rad51 (17). Armed with this knowledge, we next examined the DNA damage sensitivity of the phosphomutants in the *rad55Δ* background, where the remaining Rad51-dependent DNA repair is strictly dependent on S5S1 (6). Unlike the mild phenotypes of the *sfr1Δ* and *rad55Δ* mutants, the *rad55Δ sfr1Δ* double mutant displayed severe DNA damage sensitivity, as expected based on previous work (6, 32). Strikingly, and in stark contrast to the DNA repair proficiency observed in the presence of Rad55 **(Fig. 5A)**, both phosphomimetic mutants showed similar sensitivity to *sfr1Δ* in the *rad55Δ* background **(Fig. 5B)**. Interestingly, the *sfr1-NON* mutant also displayed a clear sensitivity to DNA damage in the *rad55Δ* background, although this was not as severe as the phosphomimetic mutants. Given that both phosphomimetic mutants phenocopied the previously established Rad51 interaction mutant, these results strongly suggest that phosphorylation serves to weaken the S5S1-Rad51 interaction and downregulate recombinational DNA repair, while also pointing towards a potentially complex regulatory mechanism in which abolishing phosphorylation is also detrimental to DNA repair.

## DISCUSSION

The potentiation of Rad51 by S5S1 is known to involve binding through two sites in the intrinsically disordered N-terminal half of Sfr1 (17). Furthermore, emerging evidence has suggested that PTMs play an important role in regulating HR (19). Here, we examined the intersection of these two phenomena. We demonstrated that phosphomimetic mutations in Sfr1 do not obviously affect the global structure of S5S1 or the disordered nature of Sfr1N **(Fig. 1)**. The physical binding of S5S1 to Rad51 was severely impaired by these mutations **(Fig. 2)**, and this translated to a clear functional defect in Rad51 filament stabilisation and stimulation of DNA strand exchange **(Figs. 3,4)**. Mutant strains of *S. pombe* expressing phosphomimetic Sfr1 showed defects in DNA repair, but only in the absence of Rad55-Rad57 **(Fig. 5)**, consistent with previous suggestions that defects in the S5S1-Rad51 interaction can be suppressed by Rad55-Rad57 (17). Importantly, the biochemical defects and DNA damage sensitivity phenotypes were comparable to a previously established Rad51 interaction mutant, providing further support for the notion that phosphorylation of Sfr1 is a critical regulator of S5S1-dependent Rad51 modulation. The implications of these findings are discussed below.

### Phosphoregulation of the S5S1-Rad51 interaction

Intrinsically disordered regions of proteins are enriched in phosphosites that often regulate their binding to interacting partners, and some phosphorylation events have even been suggested to induce folding of disordered regions (25, 26). A notable example involves the human tri-nuclease complex consisting of SLX1-SLX4, MUS81-EME, and XPF-ERCC1, which is responsible for the nucleolytic processing of a broad range of recombination and replication intermediates (33). The domain of SLX4 that binds MUS81 was shown to be disordered, and significantly, this domain underwent a disorder-to-order transition upon phosphorylation by CDK1 that enhanced its binding to MUS81 (34).

Given that the CD spectra of S5S1-PM and Sfr1N-PM were indistinguishable from wild type, phosphorylation-induced conformational changes are unlikely to be responsible for the impaired binding of S5S1-PM to Rad51. We favour a scenario where the S5S1-Rad51 interaction involves multiple electrostatic contacts, and the negative charge imparted by phosphorylation, which is mimicked by phosphomimetic mutations, disrupts these contacts. This possibility is supported by several lines of evidence. First, the interaction of Sfr1N with Rad51 displayed clear sensitivity to salt; while Sfr1N co-immunoprecipitated with Rad51 at 25 mM NaCl, this interaction was significantly diminished at 50 mM NaCl (16). Second, mutation of seven positively charged residues (K/R) to A in the two interaction sites of Sfr1N disrupted the binding of S5S1 to Rad51 (17). Third, mutation of three negatively charged residues (D/E) to A on the exterior of the Rad51 filament reduced S5S1 binding (35). We propound that positively charged sites on Sfr1 form contacts with negatively charged sites on Rad51 **(Fig. 5C)**; the addition of negatively charged phosphate groups to Sfr1 likely masks the positively charged sites and impairs binding of S5S1 to Rad51, essentially mimicking the S5S1-7A mutant in which positive residues are neutralised via mutation to A.

What kinase could be responsible for this phosphorylation? Among the five S/T residues under examination, four (T73, S109, S116, S165) match the minimal consensus sequence for CDK1 (S/T-P) (36). By contrast, S24 may be phosphorylated by Aurora kinase, another M phase regulator (37). Both S109 and S165 have been shown to be phosphorylated in M phase, when CDK1 activity is at its peak (24). Moreover, using a conditional mutant of CDK1, it was shown that the phosphorylation of S165 was significantly reduced shortly after CDK1 inhibition, with the authors concluding that this phosphorylation is likely to be direct (23). Less is known about the nature of T73 and S165 phosphorylation but given that phosphosites on the same polypeptide are more likely to be modified with similar timing (23), it is plausible that these sites are also modified by CDK1 in M phase, resulting in a phosphorylated form of S5S1 that is unable to potentiate Rad51 during M phase. This may be required to curb Rad51 activity to reduce the formation of recombination intermediates that may otherwise pose an obstacle to chromosome segregation (38, 39). In the absence of such a regulatory mechanism, genome instability would ensue, leading to increased DNA damage sensitivity, as observed for the *sfr1-NON* mutant. Alternatively, it is possible that phosphorylation of Sfr1 is required to promote the dissociation of S5S1 from Rad51 after strand invasion/exchange to prevent unproductive interactions that may hinder downstream steps of HR. Presently, it is not possible to distinguish between these two possibilities. Additional support for the involvement of CDK1 in regulating S5S1 comes from the observation that genetic interactions exist between *sfr1^+^* and the CDK activating kinase *csk1^+^*, with the authors suggesting that Csk1 may act via CDK1 to influence HR (40). Given the complex nature of these genetic interactions, it is likely that the underlying mechanisms are intricate and multi-tiered.

In addition to phosphorylation of Sfr1, *S. pombe* Rad51 is ubiquitylated by both Fbh1, an F-box family helicase, and Rrp1, a RING-domain containing SWI/SNF-family translocase (41, 42). It remains to be determined how these modifications affect Rad51 activity and their relation (if any) to Sfr1 phosphorylation. What is becoming increasing clear is that PTM of HR factors plays critical roles in modulating recombinational DNA repair (19). Mechanistically, this may be achieved via the regulation of Rad51 binding potential, as demonstrated here for S5S1.

### Interplay between S5S1 and Rad55-Rad57

In *S. pombe*, the mild sensitivity of the *rad55Δ* (or *rad57Δ*) or *swi5Δ* (or *sfr1Δ*) single mutant compared with the severe sensitivity of the double mutant — which resembles the *rad51Δ* single mutant — was interpreted to mean that Rad55-Rad57 and Swi5-Sfr1 function in independent sub-pathways of HR (6, 32). However, this was challenged by the discovery that defects in the S5S1-Rad51 interaction are suppressed by a Rad55-Rad57–dependent mechanism, and that S5S1 and Rad55-Rad57 physically interact with each other (17). Interestingly, S5S1 suppressed the defects of the *rad51-E206A* mutant, which was suggested to be specifically defective in the interaction with Rad55-Rad57 (35). Collectively, these findings led to the proposal that, although capable of functioning independently of each other, Rad55-Rad57 and S5S1 collaboratively promote Rad51-dependent DNA repair in wild-type cells (17, 35).

When combined with our biochemical analyses indicating that S5S1-PM is defective in the physical and functional interaction with Rad51, the demonstration that *sfr1-PM(D)* and *sfr1-PM(E)* phenocopy *sfr1-7A* provides further support for the existence of interplay between Rad55-Rad57 and S5S1. How Rad55-Rad57 suppresses the defects of S5S1-7A and S5S1-PM is unclear. Given that Rad55-Rad57 can interact with both Rad51 and S5S1, it was proposed that Rad55-Rad57 can function as a molecular bridge to increase the local concentration of S5S1-7A around Rad51, thereby allowing S5S1C to stimulate Rad51 (17). A reasonable inference that can be drawn from the suppression of *sfr1-NON* by Rad55 is that the DNA repair defects in this strain are also associated with a dysregulated Rad51 interaction. We hypothesise that, in the cellular context, S5S1-NON interacts with Rad51 with increased affinity compared to wild-type S5S1, which may exist in a partially phosphorylated state. This increased affinity likely leads to unproductive/untimely interactions with Rad51 that are detrimental to DNA repair (discussed above). Interestingly, budding yeast Rad55 is also phosphorylated, and this modification was shown to be important for DNA repair (43, 44). The precise biological requirement for these phosphorylation events, along with the interplay between different auxiliary factors and their PTMs, will likely be a focal point of future research.

## AUTHOR CONTRIBUTIONS

Pengtao Liang, Validation, Formal analysis, Investigation, Resources, Writing—review & editing, Visualisation

Katie Lister, Validation, Investigation, Resources, Writing—review & editing, Visualisation

Luke Yates, Conceptualisation, Methodology, Formal analysis, Investigation, Writing—review & editing, Supervision

Bilge Argunhan, Conceptualisation, Methodology, Validation, Formal analysis, Investigation, Resources, Writing—original draft, Writing—review & editing, Visualisation, Supervision, Project administration, Funding acquisition

Xiaodong Zhang, Conceptualisation, Formal analysis, Resources, Writing—original draft, Writing—review & editing, Supervision, Project administration, Funding acquisition

## ACKNOWLEDGEMENTS

We would like to thank all members of our laboratory for support and discussions, as well as Marc Morgan of the Protein Crystallography and Macromolecular Interactions Facilities for help with circular dichroism experiments. B.A. also extends his gratitude to Hiroshi Iwasaki (Tokyo Institute of Technology) for support during the early stages of the project. This work was supported by a Marie Skłodowska-Curie Actions fellowship to B.A. (101022335) and a Wellcome Trust Senior Investigator Award to X.Z. (210658/Z/18/Z).

## MATERIALS AND METHODS

### Protein purification

Rad51 and RPA were purified exactly as previously described (17). N-terminally hexahistidine-tagged Sfr1 was co-expressed with Swi5 as an operon from a pET11b vector in an *Escherichia coli* BL21 DE3 strain (NEB) that had been pre-transformed with the pRARE2 plasmid (Novagen). Plasmids used are: pBA184, S5S1; pBA186, S5S1-NON; pBA187, S5S1-PM; and pBA197, S5S1-7A. For each S5S1 variant, 2 L of *E. coli* was grown to an OD600 of 0.4–0.5, 1 mM of IPTG was added, and incubation was continued at 18°C for ∼15 h at 170 rpm. Cells were then harvested by centrifugation. All subsequent steps were carried out at 4°C. Cells (8–10 g wet weight) were resuspended in 50 mL of lysis buffer (25 mM HEPES-KOH [pH 7.5], 500 mM NaCl, 10% glycerol, 0.5 mM TCEP), lysed by sonication, and clarified by centrifugation (67,000 g 1 h). Cleared lysates were supplemented with 0.05% polyethyleneimine, incubated for 30 min with mixing, and precipitate was removed by centrifugation (25,000 g 20 min). Ammonium sulphate was slowly added to the supernatant (35% saturation), which was then incubated with mixing for 1 h. Precipitate was collected by centrifugation (10,000 g 10 min), resuspended in 50 mL of lysis buffer, and applied to a 5 mL cOmplete His-tag purification column (Roche). The column was then washed with ∼100 mL of lysis buffer and bound proteins were eluted in 25 mL of lysis buffer supplemented with 300 mM imidazole. The eluate was dialysed overnight against 1 L of H buffer (25 mM HEPES-KOH [pH 7.5], 10% glycerol, 0.5 mM TCEP) containing 200 mM KCl, diluted with 3x volumes of H buffer, and applied to a HiTrap Q column (5 mL). Proteins were eluted with a linear gradient (100–600 mM KCl, 190 mL). Peak fractions were combined, concentrated to ∼3 mL, applied to a 16/60 Superdex 200 pg gel filtration column, and developed in H buffer containing 200 mM KCl. Peak fractions were combined, concentrated, and flash-frozen in liquid nitrogen as small aliquots. Protein concentration was estimated by measuring A280 with a molar extinction coefficient of 12,490 M^-1^ cm^-1^.

Up until ammonium sulphate precipitation, N-terminally hexahistidine-tagged Sfr1N (pBA198) and Sfr1N-PM (pBA200) were treated the same as S5S1. Ammonium sulphate precipitation was at 45% saturation. Following nickel affinity purification and dialysis, the sample was applied to HiTrap Q column (5 mL) and recovered in the flow-through. The sample was then applied to a HiTrap Heparin column (5 mL) and eluted with a linear gradient (100–600 mM KCl, 100 mL). Peak fractions were concentrated and Sfr1N was further purified by gel filtration, as for Swi5-Sfr1. Protein concentration was estimated by the bicinchoninic acid assay, with wild-type S5S1 serving as a standard. Other than the His-tag purification column, all columns are from Cytiva. Chromatograms for size-exclusion chromatography are shown in Fig. 1 and are expressed relative to the highest A280 reading during each run.

### Circular dichroism spectroscopy

Global secondary structure was examined using a spectrometer (Chirascan, Applied Photophysics) with a Peltier temperature controller (Quantum Northwest). Swi5-Sfr1 or Sfr1N (wild type or mutants) were diluted to 7.35 µM in 20 mM potassium phosphate (pH 7.4) and transferred into a 1 mm path length quartz cuvette (Hellma Analytics). Circular dichroism was recorded from 190–260 nM at 20°C (0.5 nm intervals, 1 nm bandwidth, for 1 s per datapoint).

### Protein-protein interaction assays

For the in-solution pull-down assay using nickel resin **(Fig. 2A)**, 500 nM of Rad51 was mixed with 500 nM of S5S1 (wild type or mutant) in pull-down buffer (30 mM Tris-Cl [7.5], 100 mM KCl, 5 mM MgCl_2_, 1 mM ATP, 0.25 mM TCEP, 5% glycerol, 0.05% Tween-20) on ice (total volume 120 µL). 10 µL was withdrawn and mixed with SDS-PAGE loading buffer (input). The remaining reaction was incubated at 30°C for 15 min then on ice for 5 min. 10 µL of cOmplete nickel resin (Roche) was added to each reaction and hexahistidine-tagged proteins were immobilised on the resin by incubating with gentle agitation for 1 h at 4°C. The resin was pelleted by brief centrifugation and washed twice with 400 µL of pull-down buffer. Bound proteins were eluted with 40 µL of SDS-PAGE loading buffer (37°C 10 min 1300 rpm) and analysed by SDS-PAGE.

For the filament binding assay **(Fig. 2C)**, 3.33 µM Rad51 was added to filament binding buffer (25 mM Tris-Cl [7.5], 100 mM NaCl, 3.5 mM MgCl_2_, 2 mM ATP, 0.5 mM TCEP, 5% glycerol, 0.05% Tween-20) containing 10 µMnt of circular ssDNA (PhiX174 virion DNA, NEB) immobilised on magnetic Streptavidin-coated beads (Dynabeads M-280, ThermoFisher Scientific) through an oligonucleotide with a biotin moiety at its 5’ end (BA655, ATAAGGCCACGTATTTTGCAAGCTATTTAACTGGCGGCGATTGCGTACCCGACGACCAAAATTAGGGTCAACGCTACCTGTAGGAAGTGTCCGCATAAAG). Filaments were allowed to form by incubating at 37°C with gentle agitation for 5 min, then 3.33 µM S5S1 (wild type or mutant) was added, and incubation was continued as before for a further 5 min. The solution was then separated into precipitate (DNA bound) and supernatant (unbound) fractions using a magnetic stand. Proteins were eluted from the resin in 1x SDS-PAGE loading buffer (37°C 15 min 1300 rpm) and, along with the supernatant fraction, analysed by SDS-PAGE. Bound proteins were quantified in FIJI (45). Briefly, background was subtracted using the rolling ball method, then the signal corresponding to the region of the gel containing Sfr1 for the reaction where Sfr1 was omitted was subtracted from the corresponding signal for reactions containing Sfr1. Sfr1 signal was then normalised to Rad51 and plotted relative to wild type (n = 3).

### Negative stain electron microscopy

Electron microscopy buffer (25 mM HEPES-KOH [pH 7.5], 100 mM KCl, 2 mM ATP, 5 mM MgCl_2_, 0.5 mM TCEP) containing 1.5 µMnt of a 116-mer oligonucleotide (BA494, TTCAATATCTGGTTGAACGGCGTCGCGTCGTAACCCAGCTTGGTAAGTTGGATTAAGCACTCCGTGGACAGATTTGTCATTGTGAGCATTTTCATCCCGAAGTTGCGGCTCATTCT) was supplemented with Rad51 (500 nM) and incubated at 37°C for 5 min. Swi5-Sfr1 (wild type or mutants, 100 nM) was then added and incubation was continued for 10 min. 4 µL of the reaction was then applied to a carbon-coated 300 mesh copper grid (Agar Scientific) and fixed with 2% uranyl acetate. Images were captured using a Tecnai 12 electron microscope at 52,000x magnification and filament length was measured using FIJI (45). Visualisation and statistical analysis was performed in Graphpad Prism (version 8).

### DNA strand exchange assay

All concentrations indicate final concentrations in the 10 µL reaction, and all incubations were at 37°C. Strand exchange buffer (25 mM Tris-Cl [pH 7.5], 100 mM KCl, 2 mM ATP, 3.5 mM MgCl_2_, 0.5 mM TCEP, 5% glycerol, 4 mM phosphocreatine, 4 U/mL creatine kinase) containing 10 µMnt cssDNA (NEB, PhiX virion DNA) was supplemented with Rad51 (5 µM) and incubated for 8 min. The indicated concentration of Swi5-Sfr1 (wild type or mutant) was added and reactions were incubated for a further 8 min. RPA (1 µM) was then added to the reactions, and following another 8 min incubation, 10 µMnt ldsDNA (NEB, PhiX RF I digested with ApaLI) was added to initiate the strand exchange reaction. After 2 h, 1 µL of 200 µg/mL psoralen was added and the reactions were exposed to 200 µJ/cm^2^ UV (UVP, CL-1000 ultraviolet crosslinker). Reactions were then deproteinised through the addition of 2.5 µL stop solution (120 mM Tris-Cl, 60 mM EDTA, 1% SDS, 0.77 mg/mL proteinase K) and incubation for 15 min at 37°C. Reactions were separated by agarose gel electrophoresis and DNA was visualised by staining with SYBR Gold (Thermo Fisher Scientific). Images were captured using the ChemiDoc MP Imaging System (Bio-Rad). Quantification was performed in FIJI (45) by subtracting background signal (rolling ball method), then the signal corresponding to NC was expressed as a percentage of the total lane signal (sum of ldsDNA, JM divided by 1.5, NC). For JM, the signal was divided by 1.5 and then expressed as a percentage of the total lane signal.

### Electrophoretic mobility shift assay

DNA binding buffer (25 mM HEPES-KOH [pH 7.5], 100 mM KCl, 3.5 mM MgCl_2_, 0.5 mM TCEP, 5% glycerol) containing 20 nM ssDNA with a Cy5 label at the 3’ end (80-mer, BA663: TTGATAAGAGGTCATTTTTGCGGATGGCTTAGAGCTTAATTGCTGAATCTGGTGCTGTAGCTCAACATGTTTTAAATATG), or BA663 annealed to its unlabelled complementary strand (BA664), was supplemented with the denoted concentration of Swi5-Sfr1 (wild type or mutants) in a 10 µL reaction. After a 15 min incubation at 37°C, 2 µL of loading dye was added to the reaction, and samples were separated by agarose gel electrophoresis at 4°C (0.8% gel in TAE buffer). Gels were imaged using a ChemiDoc MP Imaging System (Bio-Rad). Oligonucleotides were synthesised by Eurofins and purified by HPLC.

### Spot-test to assay DNA damage sensitivity

*S. pombe* strains were prepared by tetrad dissection according to standard protocols (46) and spot-tests were conducted exactly as previously described (35). Briefly, single colonies were inoculated in rich media (YE with supplements) for 24 h, then diluted into fresh media and grown until they reached a cell density of approximately 1 x 10^7^ cells/mL. 10-fold serial dilutions were then prepared and 5 µL of cells were spotted onto solid rich media, with the most concentrated spot corresponding to 1 x 10^5^ cells. Plates were either untreated (control) or exposed to the indicated dose of ultraviolet light, then left for 3–4 days until sufficient growth was observed. Growth was always at 30°C.

---

**Figure S1.**
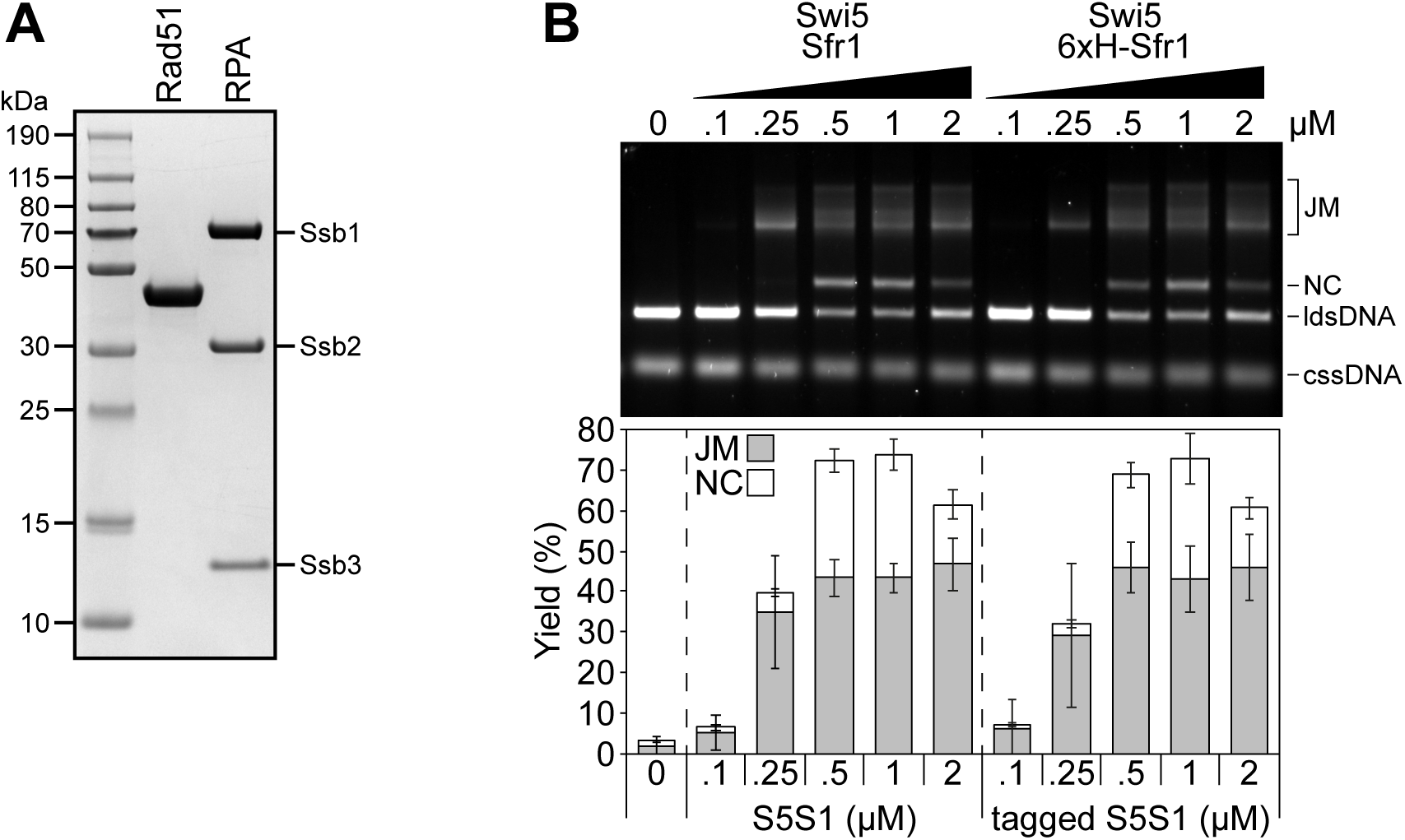
Hexahistidine-tagged S5S1 is proficient for the stimulation of Rad51-driven DNA strand exchange. **(A)** Purified Rad51 and RPA were analysed by SDS-PAGE and Coomassie staining. 2 µg of protein was loaded in each case. **(B)** Strand exchange assays were conducted as described for Fig. 4B. Averages are plotted. n = 3, error bars represent standard deviation.

**Figure S2.**
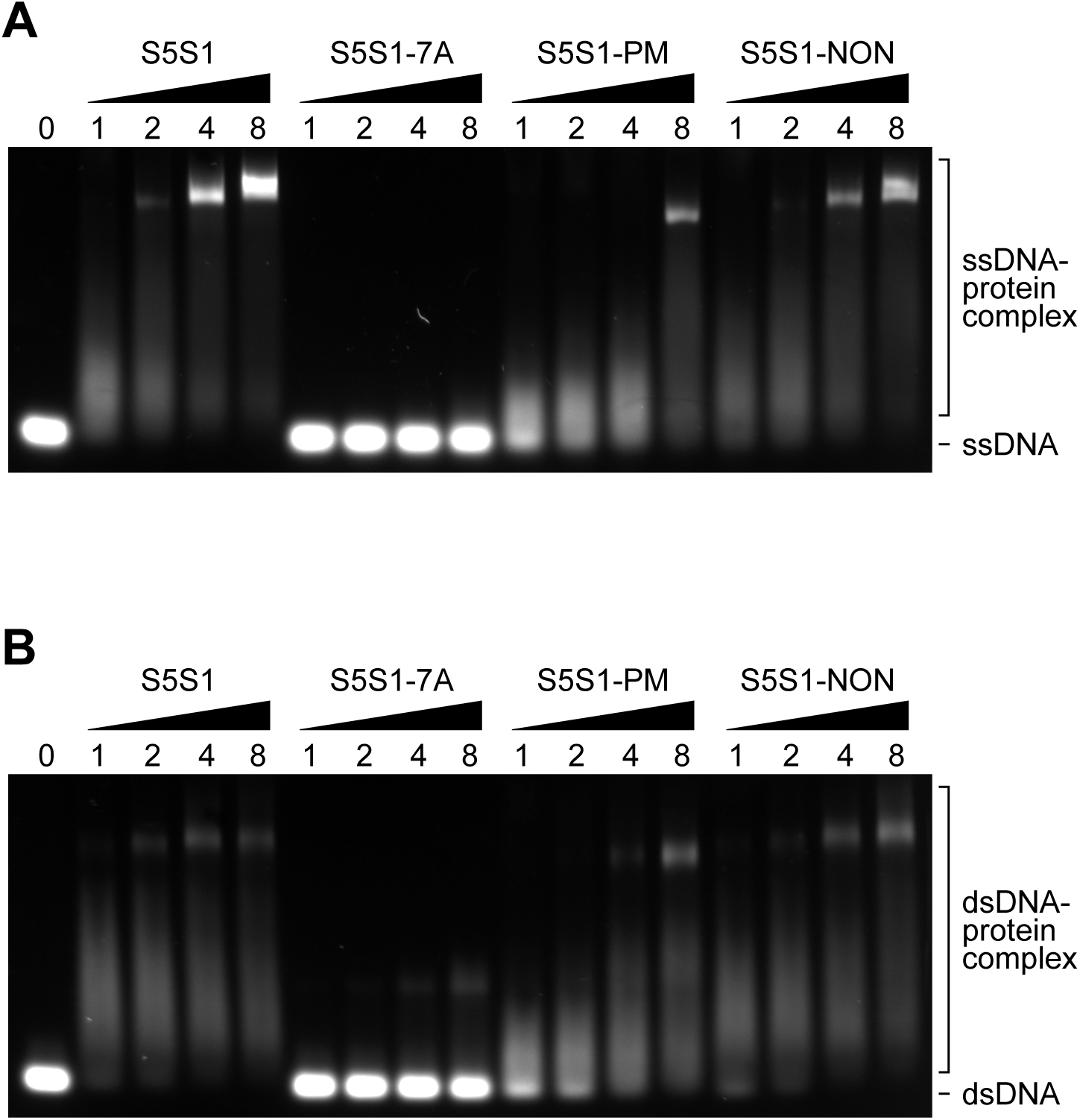
S5S1-PM is mildly impaired for DNA binding. Electrophoretic mobility shift assays were conducted with 80-mer ssDNA **(A)** or dsDNA **(B)**. DNA substrates (20 nM) were incubated with the indicated concentration of a S5S1 variant at 37°C and protein-DNA complexes were resolved by agarose gel electrophoresis.

